# Obesity/ Type II Diabetes Promotes Function-limiting Changes in Flexor Tendon Extracellular Matrix that are not Reversed by Restoring Normal Metabolic Function

**DOI:** 10.1101/143149

**Authors:** Valentina Studentsova, Keshia M. Mora, Melissa F. Glasner, Mark R. Buckley, Alayna E. Loiselle

## Abstract

Type II Diabetes (T2DM) negatively alters baseline tendon function, including decreased range of motion and mechanical properties; however, the biological mechanisms that promote diabetic tendinopathy are unknown. To facilitate identification of therapeutic targets we developed a novel murine model of diabetic tendinopathy. Mice fed a High Fat Diet (HFD) developed diet induced obesity and T2DM. Obesity/ T2DM resulted in progressive impairments in tendon gliding function and mechanical properties, relative to mice fed a Low Fat Diet (LFD), as well as a decrease in collagen fibril diameter by transmission electron microscopy. We then determined if restoration of normal metabolic function, by switching mice from HFD to LFD, is sufficient to halt the pathological changes in tendon due to obesity/T2DM. However, switching from a HFD to LFD resulted in greater impairments in tendon gliding function than mice maintained on a HFD. Mechanistically, IRβ signaling is decreased in obese/T2DM murine tendons, suggesting altered IRβ signaling as a driver of diabetic tendinopathy. However, knock-down of IRβ expression in S100a4-lineage cells (IRcKO^S100a4^) was not sufficient to induce diabetic tendinopathy as no impairments in tendon gliding function or mechanical properties were observed in IRcKO^S100a4^ relative to WT. Collectively, these data define a murine model of diabetic tendinopathy, and demonstrate that tendon-specific, rather than systemic treatment approaches are needed.

## Introduction

Tendons are dense connective tissues, composed primarily of type I collagen, that transmit forces between muscle and bone and are required for skeletal locomotion.

Type II Diabetes Mellitus (T2DM) is a metabolic disease characterized by hyperglycemia and a decrease in sensitivity to insulin (1), and is strongly associated with obesity (2). While it is difficult to dissect the systemic effects of Type II Diabetes from obesity, it is clear that induction of both obesity/T2DM leads to severe alterations in metabolic function resulting in a variety of pathological changes throughout the body (3).

While T2DM results in a plethora of systemic pathologies (4), the dramatic impact of T2DM on the musculoskeletal system has only more recently become appreciated. Approximately 83% of patients with T2DM will experience some form of musculoskeletal system degeneration or inflammation (5). Furthermore, T2DM has been shown to increase the risk of tendinopathy and tendonitis (6), with a variation in sensitivity to T2DM between different tendons (6, 7). The flexor tendons (FTs) of the hand are among the most sensitive to pathological changes of T2DM, with approximately 50% of T2DM patients experiencing limited hand function and mobility (8, 9). FTs facilitate movement of the hand and digits via nearly frictionless gliding of the FTs through the synovial sheath. Diabetic tendinopathy results in severe inflammation and pain in the hand and digits (10), and decreased digit range of motion (ROM) (6). These impairments in tendon function can severely impair use of the hands, and subsequently diminish overall quality of life. At present, the only treatments for diabetic tendinopathy are surgical release of fibrous adhesions, or corticosteroid injections to reduce local inflammation (11, 12). Consistent with disruptions in tendon homeostasis, diabetic tendons are more likely to experience spontaneous rupture requiring surgical repair (11, 12). Furthermore, tendon healing after acute injury is impaired in diabetic patients relative to non-diabetic patients (13). Therefore, there is a strong need to understand the underlying pathology of diabetic tendinopathy in order to develop therapeutic interventions to prevent disease progression.

Currently, very little is known about the cellular and molecular mechanisms underlying diabetic tendinopathy, although pathological changes increase as a function of T2DM disease duration (6). To begin to address this gap in knowledge, we have utilized a murine model of diet induced obesity and Type II diabetes (14) to test the hypothesis that flexor tendons from obese/ diabetic mice will undergo progressive, pathological changes in tendon gliding function and mechanical properties. We also examined the effects of restoration normal metabolic function as a potential means to reverse or halt diabetic tendinopathy. Finally, given the loss of insulin sensitivity that was observed in obese/ T2DM tendons, we generated tendon-specific insulin receptor conditional knockout mice to determine if IR deletion in the tendon is sufficient to recapitulate the diabetic tendinopathy phenotype.

## Results

### HFD induces Obesity and Type II Diabetes in Mice

HFD mice had a significant increase in body weight relative to LFD and HFD-LFD at all time-points, (24 weeks-LFD: 33.75g ± 2.27, HFD: 49.05g ± 2.24; HFD-LFD: 28.8g ± 7.03, p<0.0001) (Figure 1A). Body weights of HFD-LFD mice were not significantly different than LFD between 24-40 weeks, however by 48 weeks a significant increase in body weight was observed in HFD-LFD mice, relative to LFD. Fasting BG was significantly increased in HFD mice at 12, 40 and 48 weeks, relative to LFD (12 weeks-LFD: 154.6 mg/dL ± 16.1, HFD: 205.2 mg/dL ± 24.7, p <0.0001). However this increase was not observed in HFD-LFD compared to LFD between 24-48 weeks (Figure 1B). Glucose tolerance was significantly impaired in HFD, relative to LFD during a glucose tolerance test, with a 37% increase in area under the curve (AUC) in HFD, relative to LFD at 20 weeks post diet initiation (p<0.0001) (Figure 1C). In contrast, no change in glucose tolerance was observed between HFD-LFD and LFD at this time (Figure 1C). Significant increases in body fat percentage were observed in HFD, relative to LFD at 12 weeks and 48 weeks (12 weeks: +100%, p<0.0001; 48 weeks: +21%, p=0.004). Body fat percentage was significantly decreased in HFD-LFD mice, relative to both HFD (-68%, p=0.0002), and LFD (-27%, p=0.03) at 48 weeks (Figure 1D).

**Figure 1.**
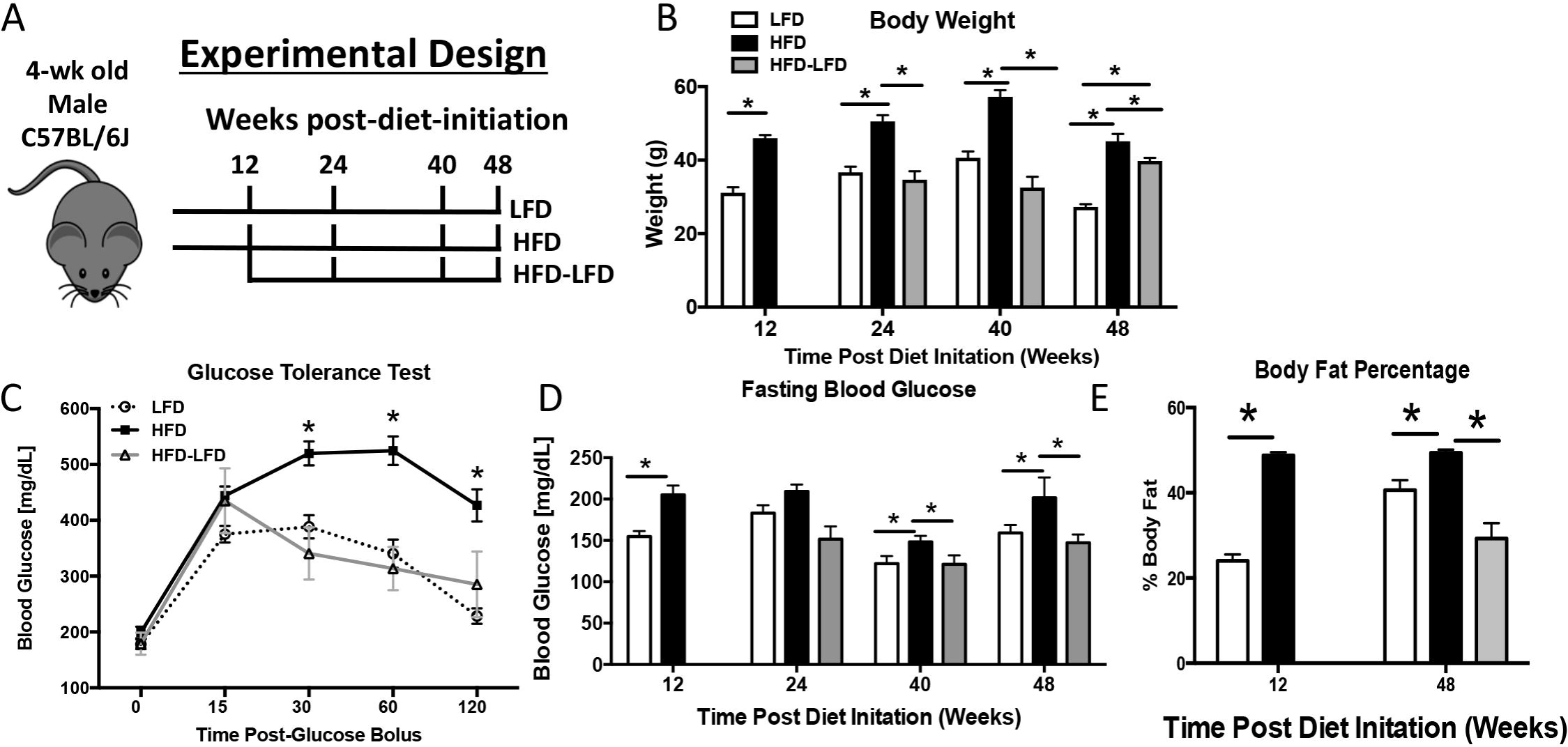
A High Fat Diet induces obesity and Type II Diabetes Mellitus. A) Male C57Bl/6J mice were initiated on either a High Fat Diet (HFD) or Low Fat Diet (LFD) at 4 weeks of age. 12 weeks after diet initiation half of the HFD cohort was randomly selected and switched to a LFD. Animals were harvested at 12, 24, 40, and 48 weeks after diet initiation (12-36 weeks after switching to LFD for the HFD-LFD cohort. B) Body weight was measured in LFD, HFD mice between 12 and 48 weeks after diet initiation. HFD-LFD mice were initiated on a HFD, and switched to a LFD 12 weeks after initiation, as such body weights from these mice were measured only between 24-48 weeks (12-36 weeks after switching to the LFD). C) At 20 weeks after diet initiation a significant impairment in glucose tolerance was observed in HFD, compared to LFD and HFD-LFD. A sustained increase in glucose levels post-glucose bolus is indicative of impaired glucose tolerance. D) Changes in fasting blood glucose were measured after a 5hr fast between 12-48 weeks after diet initiation. E) Body fat percentages of LFD, HFD, and HFD-LFD mice at 12 and 48 weeks post diet initiation. (*) Indicates p<0.05.

### HFD and HFD-LFD Impairs Tendon Range of Motion

To assess tendon range of motion we quantified metatarsophalangeal (MTP) flexion angle and gliding resistance. No change in MTP Flexion Angle was observed between groups at 12 and 24 weeks. However, at 40 and 48 weeks the MTP Flexion Angle was significantly decreased in HFD mice relative to LFD (48 weeks-LFD: 59.07° ± 5.0; HFD: 35.4° ± 9.52, p<0.0001). At 48 weeks, MTP Flexion angle was significantly decreased in HFD-LFD relative to LFD (HFD-LFD: 30.36° ± 9.3, p=0.04) (Figure 2A). Similar to MTP flexion angle between diet groups at earlier time-points, HFD and HFD-LFD at 12 and 24 weeks showed no significant changes in gliding resistance. However at 40, and 48 weeks, gliding resistance was significantly increased in HFD and HFD-LFD, relative to LFD (48 weeks-LFD: 11.4 ± 3.72; HFD: 17.48 ± 1.37; HFD-LFD: 24.64 ± 7.83, p<0.05) (Figure 2B).

**Figure 2.**
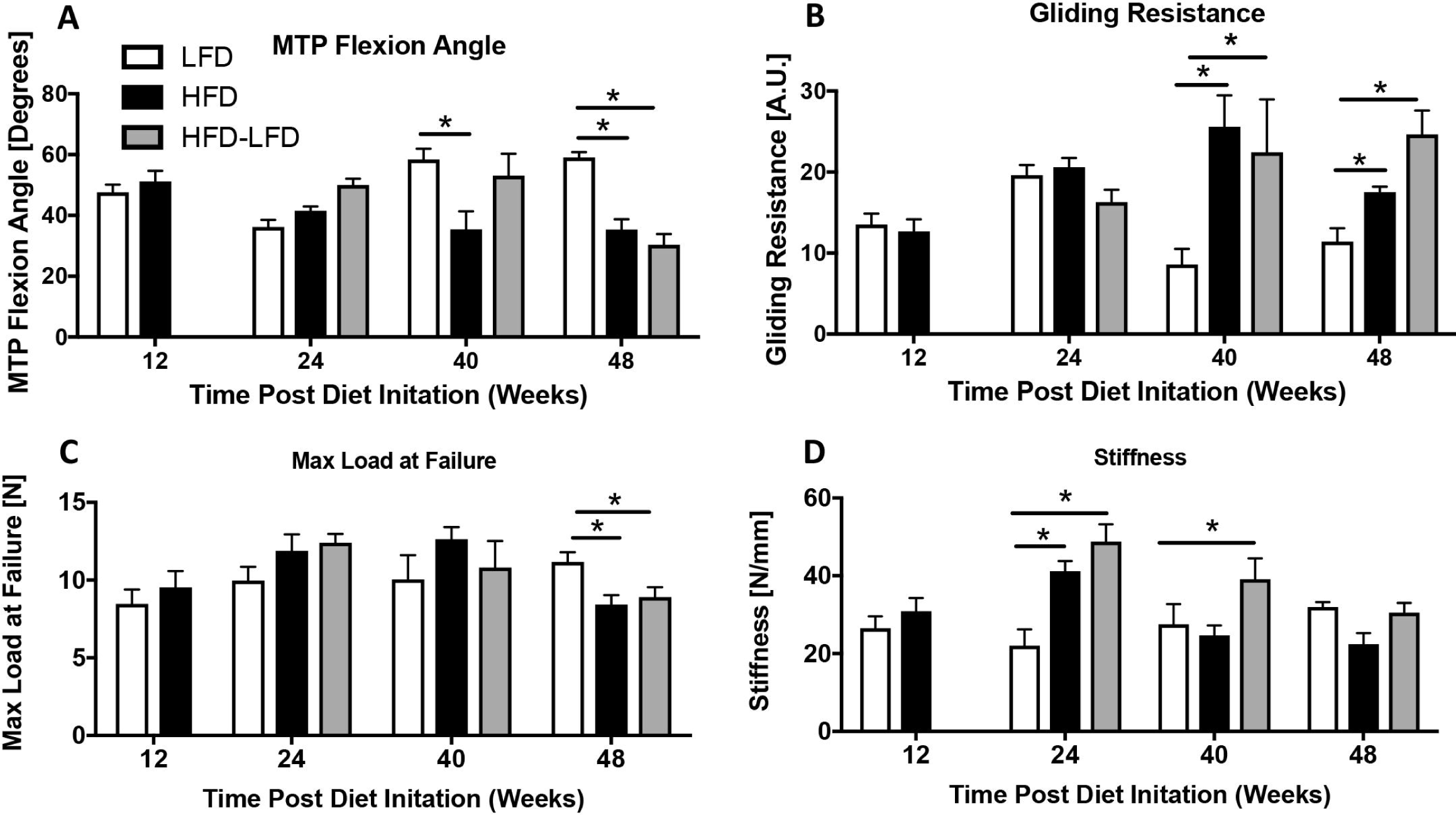
T2DM/obesity alters tendon gliding function and mechanical properties. A) Measurement of metatarsophalangeal (MTP) joint flexion angle, B) Gliding Resistance, C) Maximum load at failure, and D) Stiffness of the FDL tendon in LFD, HFD and HFD-LFD tendons between12-48 weeks after diet initiation. (*) Indicates p<0.05

### HFD and HFD-LFD alters tendon mechanical properties

Maximum load at failure was not significantly different between groups at 12, 24 and 40 weeks. At 48 weeks max load at failure was significantly decreased in HFD and HFD-LFD tendons, relative to LFD (HFD: −25%, p<0.01; HFD-LFD: −21%, p<0.04) (Figure 2C). No change in tendon stiffness was observed between groups at 12 weeks. At 24 weeks both HFD and HFD-LFD tendons exhibit a significant increase in stiffness relative to LFD (+49% and +117%, respectively, p<0.006). At 40 weeks a significant increase in stiffness was observed in HFD-LFD tendons, relative to LFD, while no differences in stiffness were observed between HFD and LFD at this time, or between any groups at 48 weeks post-diet initiation (Figure 2D).

### Obesity/ T2DM Alters Collagen Fibril Size Distribution

To define the impact of T2DM on collagen fibril density and compactness we utilized Transmission Electron Microscopy (TEM) at 40 weeks post-diet initiation. At 3500x magnification, there were no observed changes in collagen packing and organization between LFD, HFD and HFD-LFD. However, lipid deposits were observed in the mid-substance of HFD tendons (black arrows, Figure 3A). At 40,000x magnification there was a significant alteration in median collagen fibril diameter in HFD and HFD-LFD, relative to LFD (Figure 3A-C). The median fibril diameter decreased from 200.1 nm in LFD to 180.4nm in HFD (p=0.028), and 182.1 nm in HFD-LFD (p=0.046) (Figure 3B & C). No change in collagen fibril density was observed between groups at 40 weeks post-diet initiation (Figure 3D).

**Figure 3.**
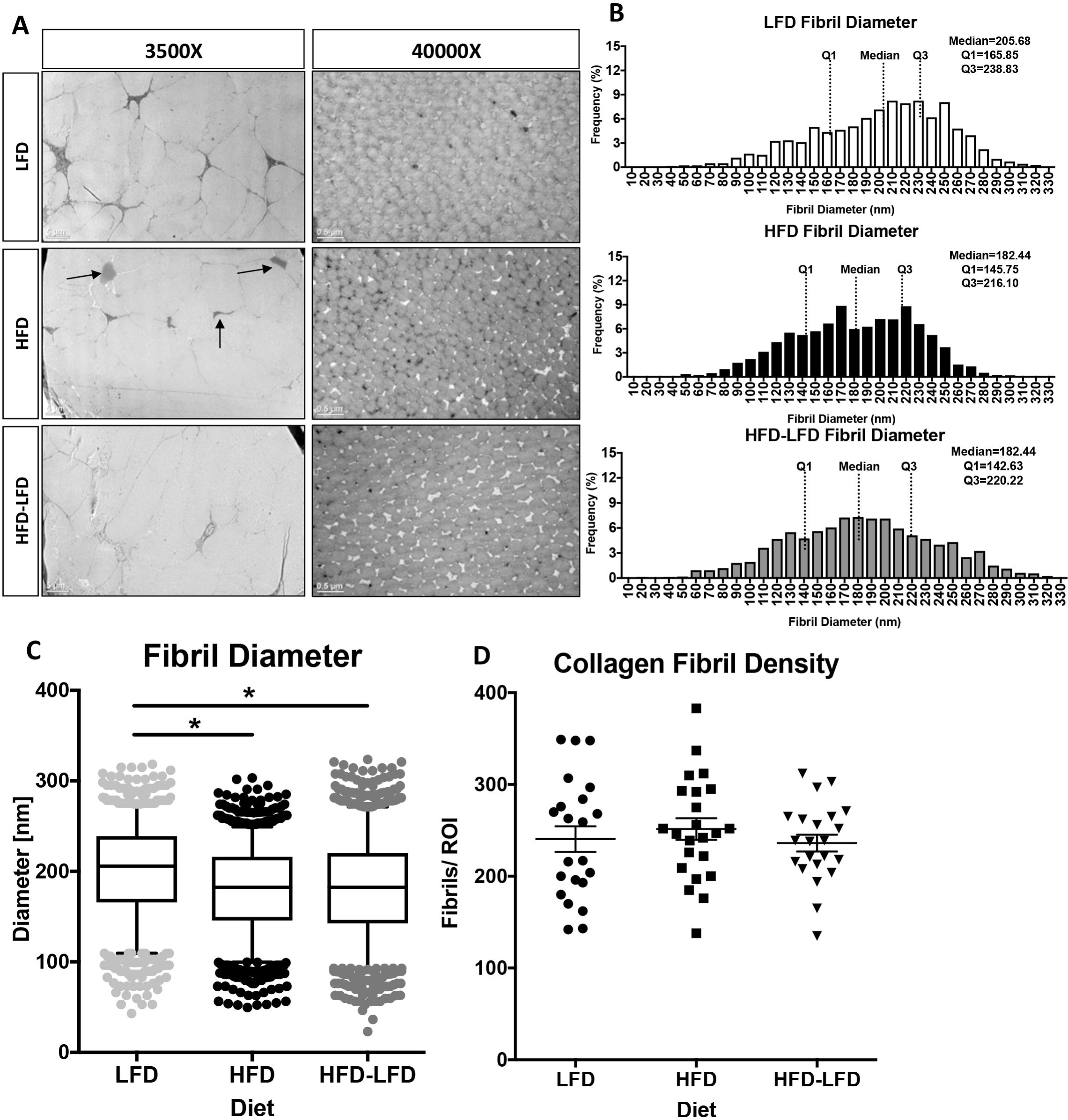
T2DM/obesity alters collagen fibril diameter. A) TEM axial images of the FDL tendon from LFD, HFD and HFD-LFD mice at 40 weeks post diet initiation. Lipid deposits in HFD tendons are noted by black arrows. Scale bars represent 5 microns in 3500X magnification images, and 0.5microns in the 40,000X magnification images. B) Collagen fibril diameter histograms demonstrating a decrease in median fibril diameter in HFD and HFD-LFD tendons compared to LFD. C) Collagen fibril diameter distributions with boxplot whiskers spanning data between the 5^th^ and 95^th^ percentiles. Data outside this range are plotted as individual points. D) Collagen fibril density. (*) Indicates p<0.05.

### Insulin Receptor Signaling is attenuated in HFD tendons

Based on changes in insulin sensitivity in other diabetic tissues (15), we examined changes in activation of Insulin Receptor (IR) signaling in primary tenocytes and obese/ T2DM and non-diabetic murine tendons. Primary tenocytes demonstrate a robust activation of IR signaling, as indicated by increased p-Akt expression, relative to vehicle treated tenocytes (Figure 4A). Upon insulin stimulation tendons from LFD mice demonstrated a robust induction of p-Akt expression, indicating activation of IR signaling. In contrast, induction of p-Akt was blunted in HFD tendons (Figure 4B).

**Figure 4.**
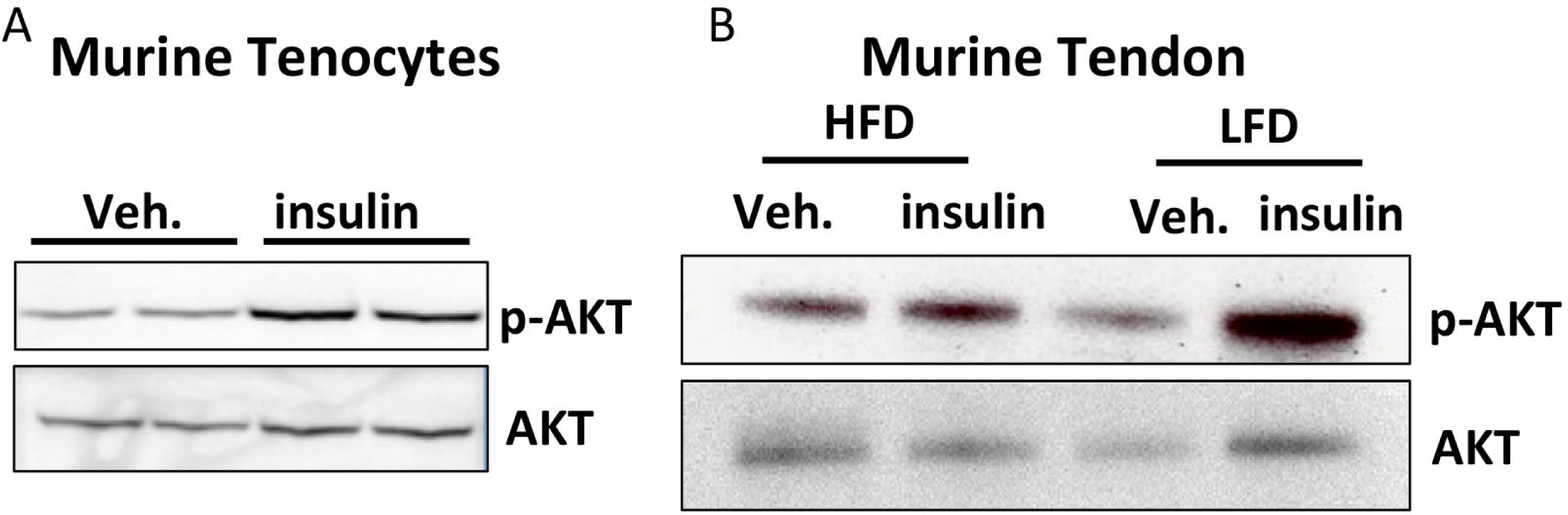
Insulin receptor expression and signaling are altered in diabetic tendons. A) Primary tenocytes isolated from non-diabetic murine tendons demonstrate strong expression of p-Akt relative to vehicle treated tenocytes, indicating activation of IR signaling in tenocytes. B) Tendons from HFD and LFD mice were stimulated with insulin or vehicle (0.5% BSA in PBS). Activation of IR signaling was observed in LFD tendons based on the increased expression of p-Akt. In contrast, no increase in p-Akt expression was observed in insulin stimulated HFD tendons compared to vehicle treatment indicating blunted sensitivity to insulin in HFD tendons.

### S100a4-Cre results in a significant decrease in insulin receptor expression in the tendon

Given the loss of insulin sensitivity in obese/T2DM murine tendons and the increase in IRβ expression in human T2DM tendons, we examined the effects of IRβ deletion in the tendon using S100a4-Cre. To demonstrate that S100a4-Cre efficiently targets resident tenocytes S100a4-Cre^+^; Ai9 reporter mice were used. Expression of Ai9 fluorescence was observed in nearly all resident tenocytes (Figure 5A). Furthermore, western blot analyses demonstrated a decrease in IRβ expression in S100a4-Cre^+^; IR^F/F^ mice (IRcKO^S100a4^), relative to wild type (WT) littermates (Figure 5B). No changes in body weight (Figure 6C), body fat percentage (Figure 5D), or fasting blood glucose (Figure 5E) were observed between WT and IRcKO^S100a4^ mice. A transient increase in glucose tolerance was observed in IRcKO^S100a4^ mice at 30 minutes after administration of the glucose bolus, however no difference in glucose tolerance was observed at 15, 60 or 120 minutes (Figure 5F). Thus, IRcKO^S100a4^ allows examination of the specific effects of IRβ knock-down in the tendon independent of obesity/ T2DM.

**Figure 5.**
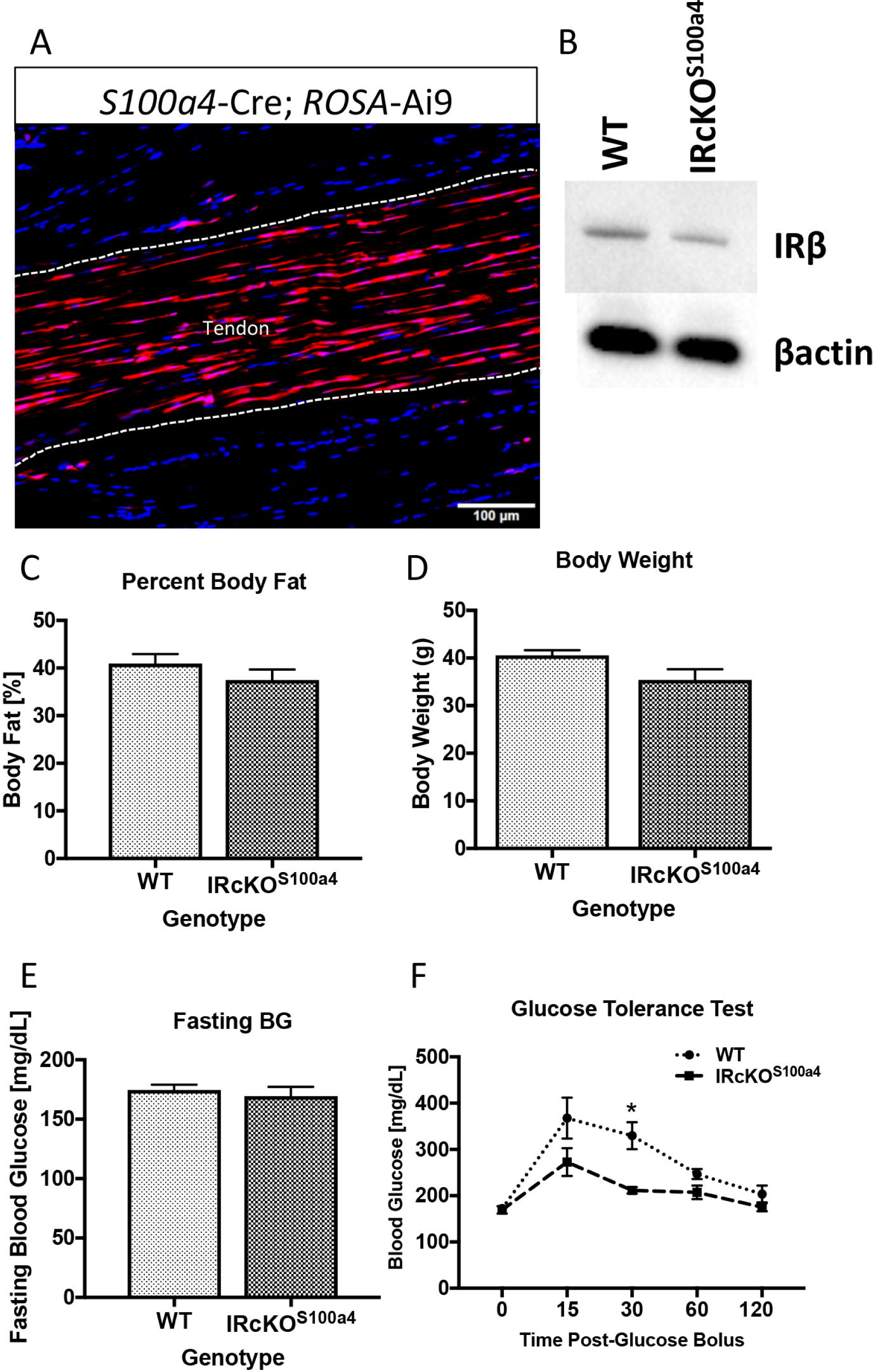
IRβ deletion in S100a4-lineage cells does not induce obesity or T2DM. A) The recombination efficiency of S100a4-Cre in the tendon was visualized using the Ai9 reporter, and demonstrates efficient recombination in the tendon. B) IRcKO^S100a4^ decreases IRβ protein expression in tendon, relative to WT. C-F) No changes in C) body weight, D) percent body fat, E) fasting blood glucose, and F) glucose tolerance were observed between WT and IRcKO^S100a4^ mice at 48 weeks of age.

**Figure 6.**
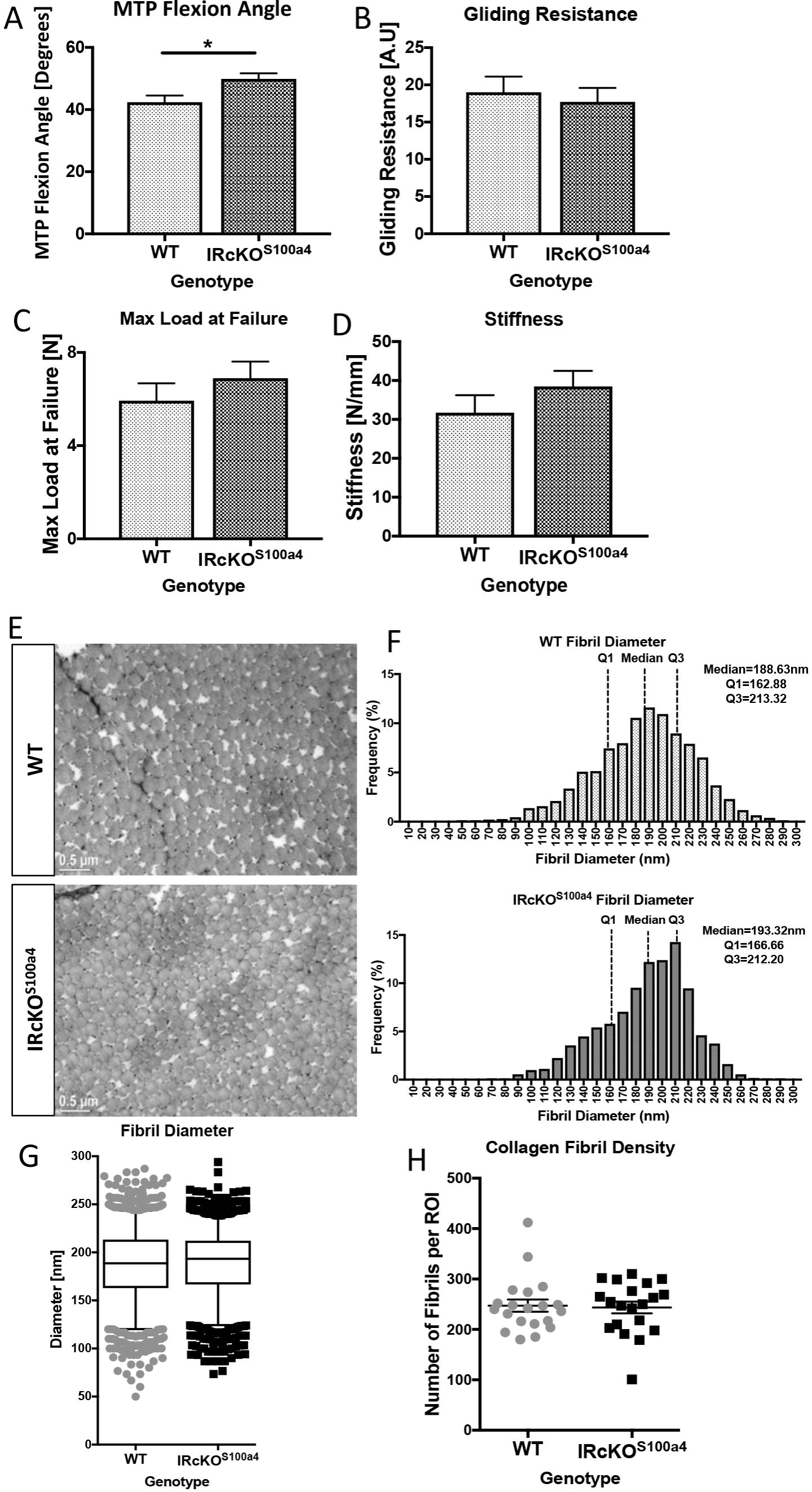
Deletion of IRβ in S100a4-lineage cells does not impair tendon gliding function, mechanical properties or collagen organization. A) MTP ROM, B) Gliding Resistance, C) Max load at failure and D) Stiffness were assessed at 48 weeks of age in WT and IRcKO^S100a4^ tendons. (*) indicates p<0.05. E) TEM axial images of the FDL tendon from WT and IRcKO^S100a4^ at 48 weeks of age. F) No changes in collagen fibril diameter were observed between WT and IRcKO^S100a4^ tendons. G) Collagen fibril diameter distributions with boxplot whiskers spanning data between the 5^th^ and 95^th^ percentiles. Data outside this range are plotted as individual points. H) Collagen fibril density of TEM images of tendons from WT and IRcKO^S100a4^ mice at 48 weeks of age.

### Insulin receptor deletion in S100a4-lineage cells does not induce diabetic tendinopathy

Tendons from IRcKO^S100a4^ had a significant increase in MTP Flexion Angle (49.87° ± 1.79, p=0.02) at 48 weeks, relative to WT (42.37° ± 2.16)(Figure 6A). No change in gliding resistance was observed between WT and IRcKO^S100a4^ (WT: 19.02 ± 2.1; IRcKO^S100a4^: 17.73 ± 1.87, p>0.05)(Figure 6B). Maximum load at failure and stiffness were not significantly different between IRcKO^S100a4^ and WT tendons (Max load: WT: 5.92N ± 0. 74, IRcKO^S100a4^: 6.9N ± 0.7, p=0.36) (Figure 6C & D). Furthermore, TEM analyses demonstrate that loss of IRβ in S100a4-lineage cells did not alter collagen fibril diameter distribution (WT Median: 188.63; IRcKO^S100a4^ Median: 193.32, p=0.21)(Figure 6E-G), or fibril density (Figure 6H)(p=0.82) at 48 weeks of age.

## Discussion

Both Type I and Type II Diabetes patients are at an increased risk of developing tendinopathy (9). While several tendons can be affected by diabetic tendinopathy, including the Achilles and the supraspinatus tendons, the flexor tendons are most commonly impacted by T2DM (8, 9). Diabetic tendinopathy decreases tendon strength and increases the likelihood of tendon rupture (11). Currently, treatment options for diabetic tendinopathy are limited to corticosteroid injections, or surgical intervention (11, 12). Therefore it is imperative to understand the biological mechanism driving this pathology in order to develop novel therapeutic approaches to improve tendon function. To address the gap in knowledge surrounding the progression of diabetic tendinopathy, we defined the effects of diet induced obesity and Type II diabetes on tendon function. To our knowledge, this is the first study to investigate the tissue-level, cellular, and molecular changes that occur during progressive diabetic tendinopathy in a murine model. We have demonstrated that induction of T2DM/obesity recapitulates the phenotypes observed clinically in diabetic flexor tendinopathy, including decreased mechanical properties and tendon range of motion (6, 11, 12). Furthermore, profound changes in collagen organization in both HFD and HFD-LFD tendons support a strong link between altered ECM organization and impaired mechanical properties in diabetic tendinopathy. Perhaps most interestingly, we demonstrate that resolution of metabolic dysfunction from obesity/T2DM is insufficient to reverse diabetic tendinopathy, highlighting the need for tendon-specific treatment options, rather than simply managing or reversing T2DM.

Clinically, one of the pathological hallmarks of diabetic tendinopathy is a decrease in tendon strength, rendering the tendon more susceptible to rupture (11, 12), while others have shown that diabetic tendons are stiffer than non-diabetic (16). Several studies have demonstrated that hyperglycemia is associated with inferior mechanical properties. Maffulli *et al.,* demonstrated that mechanical properties are reduced in the Achilles and supraspinatus tendons in a hyperglycemic environment (17), while impairments in restoration of tendon mechanical properties after injury is delayed in the Achilles tendon of hyperglycemic rats (18). Consistent with this, we observed a significant decrease in tensile strength of the FT at 48 weeks in both the HFD and HFD-LFD tendons relative to LFD, although it is not yet clear to what extent these phenotypes are driven by elevated blood glucose levels, relative to other components of metabolic dysfunction that occur in obese/ diabetic mice. Interestingly, we observed transient, significant increases in HFD (24 weeks) and HFD-LFD (24 and 40 weeks) tendons, relative LFD, consistent with clinical reports (16). However, the effects of T2DM on tendon stiffness in pre-clinical models are inconsistent. No changes were observed between HFD and LFD at 40 weeks due to a dramatic decrease in stiffness of HFD tendons between 24-40 weeks. Stiffness progressively decreased in HFD-LFD tendons with no differences observed between groups at 48 weeks. Interestingly, Connizzo *et al.,* have demonstrated a decrease in tendon stiffness in diabetic animals at the insertion, however no changes were observed in the tendon midsubstance (7), while Boivin *et al.,* observed a significant decrease in stiffness of tendons from db/db diabetic mice, relative to non-diabetic controls (19). These differential effects may be explained in part by differences in duration of T2DM, and tendon-specific differences in responsive to T2DM. Furthermore, a greater understanding of how T2DM alters the collagen ECM during the progression of diabetic tendinopathy may help explain both the changes in stiffness over time observed in this model, and the different phenotypes observed in other studies.

Tendon mechanical properties and ROM are dependent on collagen ECM organization (20). Given the decrements in mechanical properties of tendons from HFD and HFD-LFD mice, we examined changes in collagen organization and fibril diameter. TEM demonstrated significant changes in collagen fibril diameter distribution in HFD and HFD-LFD tendons, relative to LFD. These observed changes in fibril size may alter tendon gliding function, and suggests altered collagen fibril architecture as a mechanism of impaired tendon function in diabetic tendinopathy. Consistent with this, an increase in tendon bulk is observed clinically in type II diabetic patients (21), suggesting that loss of collagen compactness may impede normal gliding function leading to decreased tendon functionality and range of motion.

At the molecular level, the mechanisms driving diabetic tendinopathy are unclear. However, several studies have suggested that Advanced Glycation Endproducts (AGEs), which are dramatically increased in T2DM tissue (22), may promote loss of collagen organization due to altered cross-linking resulting from AGE-collagen binding (23). However, Fessel *et al.,* have recently demonstrated that AGEs are unable to induce tissue level impairments in tendon mechanical properties (24), suggesting other mechanisms may be driving this phenotype. To that end we have demonstrated that tendon is an insulin target tissue, and that obesity/ T2DM results in a substantial decrease in insulin sensitivity in HFD tendons at 48 weeks, suggesting that loss of IR signaling in HFD tendons may promote diabetic tendinopathy. Contrary to our hypothesis, conditional deletion of IRβ in S100a4-lineage cells did not promote changes in tendon matrix organization, gliding function or mechanical properties. However, deletion of IRβ in S100a4-lineage cells results in only a knock-down of IRβ expression, thus, we do not know what the effects of complete IRβ deletion are, or how IRβ deletion in different cell types may alter tendon structure and function. The low degree of IRβ knock-down is particularly interesting given the high efficiency we have observed in with the S100a4-Cre in other models (25) and suggests that a population other than S100a4-lineage cells are the predominant IRβ expressing cell population.

While we clearly demonstrate that a murine model of diet induced obesity and type II Diabetes recapitulates many of the clinical findings of diabetic tendinopathy, there are several limitations that must be considered. We have confined these studies to male mice, as male C57Bl/6J mice are susceptible to diet induced obesity and T2DM, while females become obese but not diabetic (26, 27). While we have shown that knockdown of IRβ in S100a4-lineage cells is insufficient to recapitulate the phenotypes observed in obese/ T2DM mice, future studies will be needed to determine if IRβ deletion, in the context of obesity, accelerates or slows tendinopathy development. We have also initiated the development of diet-induced obesity and T2DM in juvenile mice rather than adult mice, which may impact the severity or development of the phenotypes observed in our model. However, given that the incidence of obesity/T2DM is rapidly rising in the pediatric population (28), modeling this phenomenon is scientifically justified and understanding how age of onset affects diabetic tendinopathy progression is an important focus for future studies.

In summary, obesity/ T2DM is sufficient to induce an irreversible cascade of pathologic changes in the flexor tendon including altered ECM organization, ultimately leading to diminished mechanical properties and tendon ROM. Furthermore, IRβ deletion in S100a4-lineage cells does not alter tendon ECM organization or tendon function. Given that diminished insulin sensitivity is observed in obese/ T2DM tendons, future investigation in the role of IRβ in additional cell populations will be important. Finally, resolution of metabolic dysfunction was not sufficient to prevent disease progression in the tendon, highlighting the need for tendon-specific treatment options.

## Methods

### Animal Ethics

This study was carried out in strict accordance with the recommendations in the Guide for the Care and Use of Laboratory Animals of the National Institutes of Health. All animal procedures were approved by the University Committee on Animal Research (UCAR) at the University of Rochester (UCAR Number: 2014-004).

### Diet Induced Obesity and Type IIDiabetes

Male C57Bl/6J mice (Jackson Laboratories, Bar Harbor, ME) were housed in groups of five in a 12-hour light/dark cycle. At four weeks of age mice were placed on either a low fat diet (LFD; 10% Kcal, Research Diets #12450J), or high fat diet (HFD; 60% Kcal, Research Diets #D12492) for up to 48 weeks. A third cohort (HFD-LFD) of mice was placed on the HFD for 12 weeks, and then switched to the LFD until sacrifice. Mice were sacrificed at 12, 24, 40, and 48 weeks post-diet initiation, via carbon dioxide inhalation and cervical dislocation.

### Insulin Receptor conditional deletion mice

To drive loss of IRβ in the tendon, IRβ^flox/flox^ (#6955, Jackson Laboratories) (29), were crossed to *S100a4-Cre* mice (#12641, Jackson Laboratories), resulting in IRcKO^S100a4^ mice. Cre-; IRβ^flox/flox^ littermates were used as wild type (WT) controls. To confirm efficient targeting of the tendon, *S100a4-Cre* mice were crossed to Ai9 reporter mice (#7909, Jackson Laboratories) (30), resulting in tdTomato fluorescence upon Cre-mediated recombination. Mice were sacrificed at 48 weeks of age.

### Measurement of Fasting Blood Glucose

At 12, 24, 40 and 48 weeks post-diet-initiation, mice (n=6 per diet per time-point) were fasted for five hours (31), weighed, and fasting blood glucose levels were measured using a OneTouch2 blood glucometer (LifeScan Inc. Milpitas, CA). Fasting blood glucose was measured at 48 weeks of age in IRcKO^S100a4^ and WT mice (n=6 per genotype).

### Glucose Tolerance Test

To assess glucose tolerance at 20 weeks post-diet-initiation LFD, HFD and HFD-LFD mice (n=6 per diet) received a 10 uL/g bolus of 20% glucose in PBS via intraperitoneal injection following a five hour fast. Blood glucose levels were measured at 15, 30, 60 and 120 minutes after the glucose bolus. Glucose tolerance was assessed at 48 weeks of age in IRcKO^S100a4^ and WT mice.

### Assessment of Body Fat

At 12 and 48 weeks post-diet-initiation (48 weeks of age in IRcKO^S100a4^ and WT mice) the percent body fat was measured using the PIXImus dual-energy X-ray absorptiometer (DXA) (GE Lunar PIXImus, GE Healthcare, WI).

### Assessment of Gliding Function

Following sacrifice, hind limbs were harvested for gliding and biomechanical testing at 12, 24, 40, and 48 weeks post diet initiation (n=6-10 per diet per time-point.) IRcKO^S100a4^ and WT mice were harvested at 48 weeks of age (n=8-10). The FDL tendon was isolated from the myotendinous junction to the tarsal tunnel, leaving the tendon intact through the digits. The proximal end of the tendon was secured between two pieces of tape using cyanoacrylate. The tibia was secured using a custom grip and the proximal end of the FDL was incrementally loaded from 0-19 g. Upon application of each weight, an image was taken of the metatarsophalangeal (MTP) joint flexion, and measured using ImageJ (http://imagej.net). Using these images, MTP flexion angle was calculated as the difference in MTP flexion angle between unloaded (0g) and 19g. The gliding resistance was calculated by fitting the flexion data to a single-phase exponential equation. Non-linear regression was used to determine the gliding resistance (32).

### Tensile Biomechanical Testing

Following gliding function testing, the FDL was released from the tarsal tunnel, and the tibia was removed. The proximal end of the FDL and the digits were held in opposing custom grips in an Instron device (Instron 8841 DynaMight axial servohydraulic testing system, Instron Corporation, Norwood, MA). The tendon was displaced in tension at 30mm/minute until failure. Stiffness and maximum load at failure were calculated from the force displacement curve.

### Transmission Electron Microscopy

FDL tendons were isolated (n=3 per diet or genotype) and fixed in Glutaraldehyde Sodium Cacodylate fixative. One-micron axial sections were cut and stained with Toluidine blue. One-micron sections were then trimmed to 70nm and stained with uranyl acetate and lead citrate. Sections were placed on grids for imaging on a Hitachi 7650 Analytical TEM. Eight to twelve nonoverlapping images were taken from mid-substance of each tendon at 15,000x and 40,000x magnification. For measurement of fibril diameter, a region of interest (ROI) of was determined within each image so that a minimum of 80 fibrils could be measured. Diameters were measured along the y-axis. Collagen Fibril density was measured in a 2340 x 1860 pixel area. Collagen fibril density and diameter measurements were made in ImageJ.

### Insulin receptor expression and signaling

Primary tenocytes were isolated from C57Bl/6J mice and used up to passage 3. Whole FTs were isolated from HFD and LFD-fed mice 48 weeks after diet initiation (n=3 per group). Tenocytes and tendons were stimulated with either vehicle (0.5% BSA in PBS) or 1nM insulin for 15 minutes, followed by protein isolation for western blot analyses. Blots were probed for phospho-AKT Ser473 (1:1000, Cell Signaling, #4060), and total AKT (1:1000, Cell Signaling, #9272S) and were developed with SuperSignal West Pico or Femto Chemiluminescent Substrate and imaged on a GelDocXR (BioRad, Hercules, CA).

### Statistical Analyses

For obesity/T2DM studies, body weight, blood glucose levels, biomechanical and gliding data were analyzed using a two-way analysis of variance (ANOVA) followed by Bonferroni's multiple comparisons with significance set at α = 0.05. TEM and body fat percentage were analyzed using one-way ANOVA with Bonferroni’s post-hoc multiple comparisons. Statistical differences between WT and IRcKO^S100a4^ were determined using un-paired t-tests. All data were analyzed using Prism GraphPad 7.0 statistical software. Data are presented as mean ± SEM.

## Acknowledgements

We would like to thank the Histology, Biochemistry and Molecular Imaging (HBMI) Core and Biomechanics, Biomaterials and Multimodal Tissue Imaging (BBMTI) Core in the Center for Musculoskeletal Research, the Multiphoton Imaging Core, and Electron Microscopy Core at the University of Rochester Medical Center for technical assistance.

This work was supported by NIAMS/NIH grants K01AR068386 (AEL), P30 AR061307 Pilot (AEL), a University of Rochester University Research Award (AEL & MRB), R01AR070765 (MRB) and R01AR056696. The Histology, Biochemistry and Molecular Imaging (HBMI) Core and Biomechanics and Multimodal Tissue Imaging (BMTI) Core are supported in part by P30 AR069655.The content is solely the responsibility of the authors and does not necessarily represent the official views of the National Institutes of Health.

## Author contributions

Study conception and design: AEL; Acquisition of data: VS, MG, KM; Analysis and interpretation of data: VS, KM, MRB, AEL; Drafting of manuscript: MG, AEL; Revision and approval of manuscript: VS, KM, MG, MRB, AEL.

## Data Availability Statement

All data generated or analyzed during this study are included in this article.

## References

1. Williamson RT. Causes of diabetes. 1909. Practitioner. 2009;253(1718):37.

2. Mokdad AH, Ford ES, Bowman BA, Dietz WH, Vinicor F, Bales VS, et al. Prevalence of obesity, diabetes, and obesity-related health risk factors, 2001. Jama. 2003;289(1):76-9.

3. Guh DP, Zhang W, Bansback N, Amarsi Z, Birmingham CL, Anis AH. The incidence of co-morbidities related to obesity and overweight: a systematic review and meta-analysis. BMC Public Health. 2009;9:88.

4. Rodriguez A, Delgado-Cohen H, Reviriego J, Serrano-Rios M. Risk factors associated with metabolic syndrome in type 2 diabetes mellitus patients according to World Health Organization, Third Report National Cholesterol Education Program, and International Diabetes Federation definitions. Diabetes Metab Syndr Obes. 2011;4:1–4.

5. Douloumpakas I, Pyrpasopoulou A, Triantafyllou A, Sampanis C, Aslanidis S. Prevalence of musculoskeletal disorders in patients with type 2 diabetes mellitus: a pilot study. Hippokratia. 2007;11(4):216-8.

6. Abate M, Schiavone C, Salini V, Andia I. Occurrence of tendon pathologies in metabolic disorders. Rheumatology (Oxford). 2013;52(4):599–608.

7. Connizzo BK, Bhatt PR, Liechty KW, Soslowsky LJ. Diabetes alters mechanical properties and collagen fiber re-alignment in multiple mouse tendons. Annals of biomedical engineering. 2014;42(9):1880–8.

8. Singla R, Gupta Y, Kalra S. Musculoskeletal effects of diabetes mellitus. J Pak Med Assoc. 2015;65(9):1024–7.

9. Wyatt LH, Ferrance RJ. The musculoskeletal effects of diabetes mellitus. J Can Chiropr Assoc. 2006;50(1):43–50.

10. Amadio PC. Friction of the gliding surface. Implications for tendon surgery and rehabilitation. J Hand Ther. 2005;18(2):112–9.

11. Tozer S, Duprez D. Tendon and ligament: development, repair and disease. Birth Defects Res C Embryo Today. 2005;75(3):226–36.

12. Sharma P, Maffulli N. Tendon injury and tendinopathy: healing and repair. J Bone Joint Surg Am. 2005;87(1):187–202.

13. Ahmed AS, Schizas N, Li J, Ahmed M, Ostenson CG, Salo P, et al. Type 2 diabetes impairs tendon repair after injury in a rat model. J Appl Physiol (1985). 2012; 113(11): 1784–91.

14. Surwit RS, Feinglos MN, Rodin J, Sutherland A, Petro AE, Opara EC, et al. Differential effects of fat and sucrose on the development of obesity and diabetes in C57BL/6J and A/J mice. Metabolism. 1995;44(5):645–51.

15. Kang L, Ayala JE, Lee-Young RS, Zhang Z, James FD, Neufer PD, et al. Diet-induced muscle insulin resistance is associated with extracellular matrix remodeling and interaction with integrin alpha2beta1 in mice. Diabetes. 2011;60(2):416–26.

16. Gefen A, Megido-Ravid M, Azariah M, Itzchak Y, Arcan M. Integration of plantar soft tissue stiffness measurements in routine MRI of the diabetic foot. Clinical biomechanics. 2001;16(10):921–5.

17. Maffulli N, Longo UG, Maffulli GD, Khanna A, Denaro V. Achilles tendon ruptures in diabetic patients. Arch Orthop Trauma Surg. 2011;131(1):33–8.

18. Korntner S, Kunkel N, Lehner C, Gehwolf R, Wagner A, Augat P, et al. A high-glucose diet affects Achilles tendon healing in rats. Sci Rep. 2017;7(1):780.

19. Boivin GP, Elenes EY, Schultze AK, Chodavarapu H, Hunter SA, Elased KM. Biomechanical properties and histology of db/db diabetic mouse Achilles tendon. Muscles Ligaments Tendons J. 2014;4(3):280–4.

20. Screen HR, Berk DE, Kadler KE, Ramirez F, Young MF. Tendon functional extracellular matrix. J Orthop Res. 2015;33(6):793–9.

21. Akturk M, Karaahmetoglu S, Kacar M, Muftuoglu O. Thickness of the supraspinatus and biceps tendons in diabetic patients. Diabetes Care. 2002;25(2):408.

22. Giacco F, Brownlee M. Oxidative stress and diabetic complications. Circulation research. 2010;107(9):1058–70.

23. Eriksen C, Svensson RB, Scheijen J, Hag AM, Schalkwijk C, Praet SF, et al. Systemic stiffening of mouse tail tendon is related to dietary advanced glycation end products but not high-fat diet or cholesterol. J Appl Physiol (1985). 2014;117(8):840–7.

24. Fessel G, Li Y, Diederich V, Guizar-Sicairos M, Schneider P, Sell DR, et al. Advanced glycation end-products reduce collagen molecular sliding to affect collagen fibril damage mechanisms but not stiffness. PloS one. 2014;9(11):e110948.

25. Ackerman JE, Best KT, O’Keefe RJ, Loiselle AE. Deletion of EP4 in S100a4-lineage cells reduces scar tissue formation during early but not later stages of tendon healing. Sci Rep. 2017;7(1):8658.

26. Surwit RS, Kuhn CM, Cochrane C, McCubbin JA, Feinglos MN. Diet-induced type II diabetes in C57BL/6J mice. Diabetes. 1988;37(9):1163–7.

27. Pettersson US, Walden TB, Carlsson PO, Jansson L, Phillipson M. Female mice are protected against high-fat diet induced metabolic syndrome and increase the regulatory T cell population in adipose tissue. PloS one. 2012;7(9):e46057.

28. D’Adamo E, Caprio S. Type 2 diabetes in youth: epidemiology and pathophysiology. Diabetes Care. 2011;34 Suppl 2:S161–5.

29. Bruning JC, Michael MD, Winnay JN, Hayashi T, Horsch D, Accili D, et al. A muscle-specific insulin receptor knockout exhibits features of the metabolic syndrome of NIDDM without altering glucose tolerance. Mol Cell. 1998;2(5):559–69.

30. Madisen L, Zwingman TA, Sunkin SM, Oh SW, Zariwala HA, Gu H, et al. A robust and high-throughput Cre reporting and characterization system for the whole mouse brain. Nat Neurosci. 2010; 13(1):133–40.

31. Ayala JE, Samuel VT, Morton GJ, Obici S, Croniger CM, Shulman GI, et al. Standard operating procedures for describing and performing metabolic tests of glucose homeostasis in mice. Dis Model Mech. 2010;3(9-10):525–34.

32. Hasslund S, Jacobson JA, Dadali T, Basile P, Ulrich-Vinther M, Soballe K, et al. Adhesions in a murine flexor tendon graft model: Autograft versus allograft reconstruction. J Orthop Res. 2008;26(6):824–33.

